# Repeat expansions in *C9orf72* rewire the 3D chromatin landscape in ALS

**DOI:** 10.64898/2026.02.09.704902

**Authors:** Tatiana Ulloa Avila, Jingying Wang, Lydia Adams, Ellen Hu, Alvaro A. Beltran, Alexey Kozlenkov, Shreyan Urhekar, Ariana B. Marquez Gonzalez, Hun-Goo Lee, J. Mauro Calabrese, Stella Dracheva, Adriana S. Beltran, Won Mah, Hyejung Won

**Affiliations:** Neuroscience Center, University of North Carolina at Chapel Hill, Chapel Hill, NC, USA; Department of Genetics, University of North Carolina at Chapel Hill, Chapel Hill, NC, USA; Department of Biostatistics, University of North Carolina at Chapel Hill, Chapel Hill, NC, USA; Friedman Brain Institute, Icahn School of Medicine at Mount Sinai, New York, NY, USA; Department of Psychiatry, Icahn School of Medicine at Mount Sinai, New York, NY, USA; Research & Development and VISN2 MIREC, James J. Peters VA Medical Center, Bronx, NY, USA; Department of Genetic Medicine, School of Medicine, Johns Hopkins University, Baltimore, MD, USA; Advanced Academic Programs, Krieger School of Arts and Sciences, Johns Hopkins University, Baltimore, MD, USA; Department of Pharmacology, University of North Carolina, Chapel Hill, NC, USA; RNA Discovery Center, University of North Carolina, Chapel Hill, NC, USA; Lineberger Comprehensive Cancer Center, University of North Carolina, Chapel Hill, NC, USA; Human Pluripotent Cell Core, University of North Carolina, Chapel Hill, NC, USA

## Abstract

Amyotrophic lateral sclerosis (ALS) is frequently driven by GGGGCC short tandem repeat (STR) expansions in *C9orf72*, yet the mechanisms by which these expansions lead to neurodegeneration remain incompletely understood. Here, we propose a novel mechanism involving higher-order chromatin architecture where *C9orf72*-STR expansions induce widespread, neuron-specific gains in chromatin loops that are closely linked to transcriptomic dysregulation in ALS. These ectopic loops colocalize with the genomic binding sites of *C9orf72*-STR RNAs and the architectural protein CTCF, supporting a model in which RNA-DNA interactions promote aberrant loop formation. Together, our findings demonstrate how *C9orf72*-STR expansions remodel the neuronal genome and disrupt gene expression, uncovering an RNA-driven mechanism of chromatin reorganization in C9-ALS that connects altered nuclear topology to gene dysregulation in neurodegeneration.

## Introduction

Amyotrophic lateral sclerosis (ALS) is a fatal and intractable neurodegenerative disorder characterized by progressive motor neuron degeneration and muscle weakness, ultimately leading to respiratory failure and death. Although most ALS cases are sporadic, the largest proportion (∼40%) of familial ALS has been linked to a single genetic driver: the pathogenic expansion of GGGGCC (G4C2) short tandem repeats (STRs) in the first intron of the *C9orf72* gene^1,2^. Despite this well-established genetic link, developing disease-modifying therapies for *C9orf72*-mediated ALS (C9-ALS) remains challenging, largely due to the limited understanding of the molecular mechanisms by which STR expansions drive disease pathology.

The *C9orf72* STR (C9-STR) expansion is thought to contribute to ALS through various different mechanisms, including *C9orf72* haploinsufficiency, the generation of sense and antisense transcripts which form RNA foci that sequester RNA-binding proteins (RBPs), and the translation of dipeptide repeat proteins (DPRs) that aggregate into toxic inclusions^3,4^. Recently, a growing body of research suggests that aberrant RNA species derived from C9-STRs may induce pathological rewiring of chromatin architecture. C9-STR RNAs can form multimolecular G-quadruplexes (mG4s)^5,6^, which have been shown to promote and stabilize the formation of RNA-DNA hybrids (R-loops) that strengthen CTCF binding and facilitate chromatin looping^7,8^. Beyond C9-STRs, other disease-associated STRs colocalize with boundaries of topologically associating domains (TADs), suggesting that STRs may disrupt TAD organization and thereby local chromatin structure^9^. In support of this hypothesis, the pathogenic STRs in the *FMR1* gene, which cause fragile X syndrome (FXS), have been shown to disrupt the local TAD boundaries and alter global inter-chromosomal interactions^10^. These findings implicate disease-associated STRs as potential architects of pathogenic chromatin rewiring.

Here, we investigate the impact of C9-STRs on chromatin architecture by generating genome-wide chromosome conformation maps from neurons and glia isolated from postmortem cortical tissue of C9-ALS patients and neurotypical controls, as well as from hiPSC-derived neurons with and without the C9-STR expansion. While we did not observe disruptions in local chromatin structure at the *C9orf72* locus, we detected a genome-wide increase in loop formation that was closely associated with repression of neuronal genes in C9-ALS. This ectopic loop formation was evident in both postmortem neurons from C9-ALS patients and in hiPSC-derived neurons carrying the C9-STR expansion, but not in glial cells. Building on prior evidence linking noncanonical RNA structures to chromatin organization, we mapped the genomic binding sites of C9-STR RNAs to examine RNA-DNA interactions. These RNA-bound regions were notably enriched for both ectopic chromatin loops and CTCF binding sites. This suggests a potential mechanistic interplay between repeat-derived RNA, architectural proteins, and chromatin topology, offering new insight into how structural disorganization may contribute to ALS pathogenesis.

### Global chromatin reorganization in C9-ALS

To understand the impact of C9-STR expansion on 3D chromatin structure, we generated Hi-C libraries from neurons (NeuN-positive cells) and glia (NeuN-negative cells) isolated from the cortex of C9-ALS and neurotypical controls (**Figure 1A**, **Methods**).

**Figure 1.**
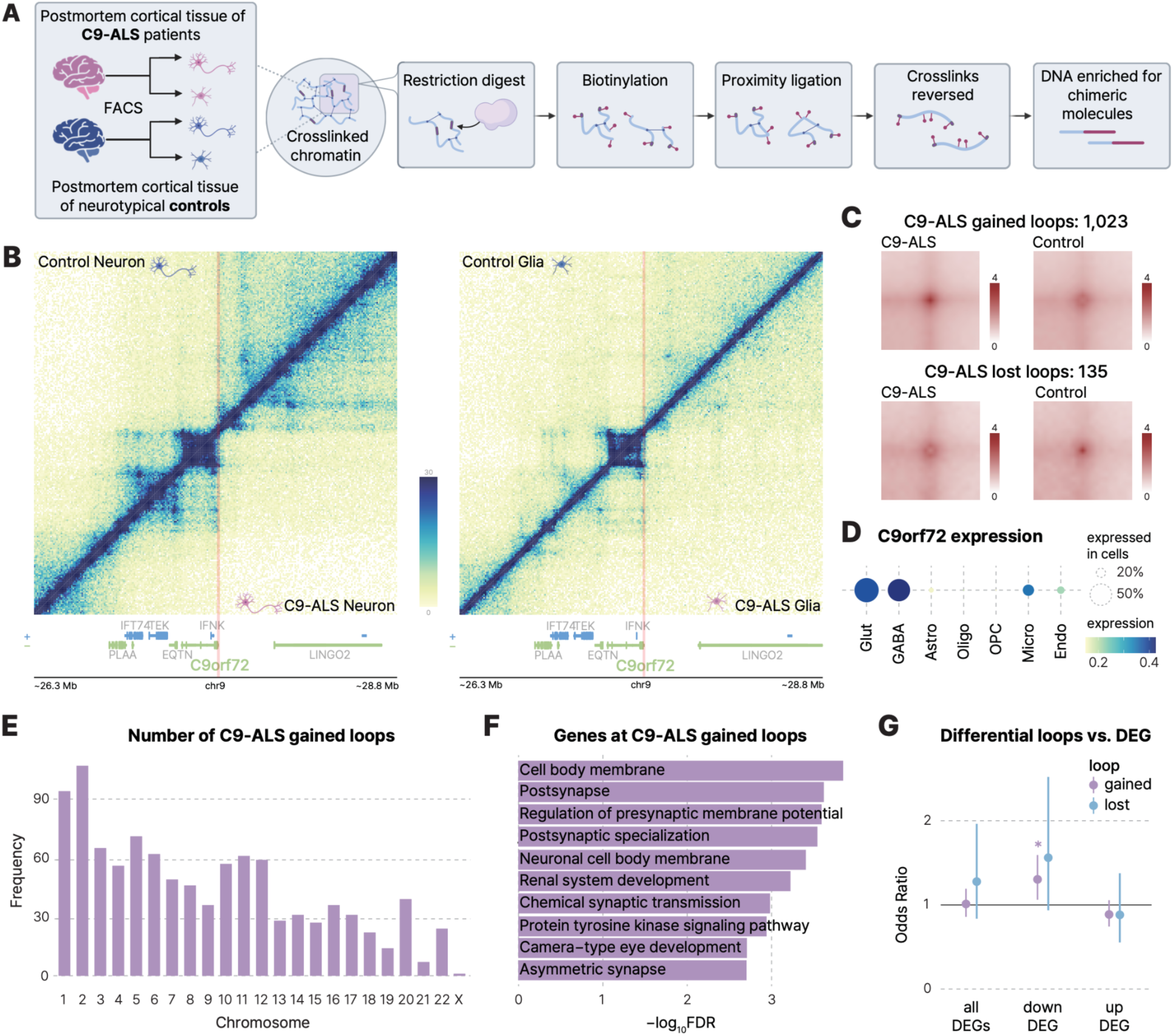
Global chromatin rewiring in C9-ALS neurons. **A.** Genome-wide chromosome conformation maps were generated from neurons and glia isolated from postmortem brain samples of C9-ALS patients and neurotypical controls. **B.** Local chromatin interaction at the *C9orf72* locus in neurons (left) and glia (right). Chromatin organization in neurotypical controls is shown in the top left, while chromatin organization in C9-ALS is shown in the bottom right. **C.** Aggregated Peak Analysis (APA) results for C9-ALS gained loops and C9-ALS lost loops in neurons. **D.** Cell type-specific expression of *C9orf72* in the cortex. Glut, glutamatergic neurons; GABA, GABAergic neurons; Astro, astrocytes; Oligo, oligodendrocytes; OPC, oligodendrocyte precursor cells; Micro, microglia; Endo, endothelial cells. **E.** Distribution of C9-ALS gained neuronal loops across chromosomes, showing an even genome-wide spread. **F.** Gene ontology analysis of genes anchored at C9-ALS gained loops. **G.** C9-ALS gained loops are enriched for downregulated genes in C9-ALS. DEG, differentially expressed genes. Two-sided Fisher’s exact test, *FDR<0.05.

We first assessed whether the local chromatin structure of the *C9orf72* locus was disrupted in C9-ALS. Although C9-STRs are located at TAD boundaries^9^, we did not observe a clear disruption of local TADs in C9-ALS, in either neurons or glia (**Figure 1B**). This result suggests that the presence of the C9-STR expansion does not necessarily lead to local chromatin structure disruption. However, the *C9orf72* locus exhibited cell type-specific chromatin architecture, with glia displaying a more dispersed TAD structure compared to the distinct TAD structure observed in neurons (**Figure 1B**).

Since the C9-STR expansion did not disrupt local chromatin structure, we next examined global chromatin organization in C9-ALS. Interchromosomal interactions remained largely unaffected in both neurons and glia from C9-ALS patients compared to neurotypical controls (**Figure S1**).

In line with minimal large-scale architectural reorganization, A/B compartment switching was detected in only 6% of the genome in neurons and 9% in glia in C9-ALS compared to neurotypical controls (**Figure S2A-B, Table S1**). To further evaluate compartment changes consistently observed in C9-ALS, we performed a differential compartment analysis^11^. Among genomic regions with significant compartment changes between C9-ALS neurons and controls, a greater proportion (27.63%) transitioned from compartment B to A, while a smaller fraction (12.06%) transitioned from A to B, suggesting a global shift toward more transcriptionally active compartments in C9-ALS neurons (**Figure S2A, C**). In contrast, C9-ALS glia did not exhibit a similar imbalance in compartment switching, with 30.92% of differential regions transitioning from A to B and 34.20% from B to A (**Figure S2B, D**).

Given the established relationship between chromatin compartments and gene expression^12^, we next evaluated whether regions undergoing compartment switching harbor genes that are differentially expressed in C9-ALS patients^13^. However, we did not observe a significant relationship between differential compartments and previously reported transcriptomic alterations in C9-ALS, in either neurons or glia (**Figure S2E-F**)^13–15^, suggesting that this level of 3D genome reorganization may be uncoupled from molecular neuropathology in C9-ALS.

We next examined alterations in chromatin looping in C9-ALS, given their central role in organizing the 3D genome by mediating enhancer-promoter interactions and insulation of regulatory domains^16–18^. Unexpectedly, we observed a ∼40% increase in the number of chromatin loops in C9-ALS neurons, but not in glia (**Figure S3A**). Since the number of detected loops can be influenced by various technical confounders, we performed a differential loop analysis^19^ between neurons from C9-ALS patients and neurotypical controls (**Methods**, **Table S2**). We identified 1,023 loops that were significantly gained in C9-ALS neurons compared to neurotypical controls (hereafter referred to as C9-ALS gained loops) and 135 loops that were significantly lost (hereafter referred to as C9-ALS lost loops, **Figure 1C**). This ∼10-fold excess of gained over lost loops mirrors the global increase in chromatin looping observed in C9-ALS neurons (**Figure S3A**). In contrast, no such imbalance in differential looping was observed in C9-ALS glia relative to controls (**Figure S3B**), highlighting a neuron-specific chromatin reorganization. This specificity may be attributable to higher expression levels of *C9orf72* in cortical neurons compared to glia^20^ (**Figure 1D**). C9-ALS gained loops were distributed broadly across the genome (**Figure 1E**), suggesting that the C9-STR expansion drives widespread reorganization of 3D genome architecture.

To assess whether these gained loops are associated with the emergence of new TADs, we identified C9-ALS associated TAD reorganization using DiffDomain^21^ (**Table S3**). Consistent with the increase in C9-ALS gained loops, we found that TAD splitting events were substantially more frequent than TAD merging events in C9-ALS neurons (**Figure S3C**). Although C9-ALS glia also exhibited more TAD splitting than merging events (**Figure S3C**), this imbalance was markedly more pronounced in neurons (**Figure S3C,** Fisher’s exact test, p=0.042). Notably, 30.6% of TAD splitting events in neurons overlapped with C9-ALS gained loops, compared to only 10.3% in glia, suggesting that ectopic loop formation may be accompanied by local domain reorganization (**Figure S3D**).

We next explored which genes might be impacted by the C9-ALS gained and lost loops by examining the genes whose promoters were anchored to the differential loops. Genes linked to C9-ALS gained loops were enriched for synaptic and neuronal functions, suggesting that these ectopic loops may contribute to altered neuronal function and degeneration in C9-ALS (**Figure 1F**). In contrast, C9-ALS lost loops were associated with genes enriched for peroxisomal and microbody membrane functions (**Figure S3E**).

Given the established connection between chromatin looping and gene regulation^22,23^, we investigated whether differential loops in C9-ALS were associated with transcriptomic alterations previously reported in C9-ALS postmortem brain samples^13^. While no clear relationship was observed between transcriptomic dysregulation and C9-ALS lost loops—likely due to their small number—promoters of genes downregulated in C9-ALS neurons were more frequently anchored to gained loop anchors compared to stable loops (i.e., loops that did not differ between C9-ALS and neurotypical controls, **Figure 1G**). Together, these results suggest that STR-mediated 3D genome reorganization may contribute to the widespread gene expression changes observed in C9-ALS^13–15^.

### Ectopic loop formation in hiPSC-derived neurons with C9-STR expansion

Because neurons and glia isolated from postmortem brain tissue capture the endpoint rather than the onset of disease (**Figure 1A**), it remains unclear whether the observed gain in chromatin loops reflects a causal mechanism or a downstream consequence of C9-ALS pathogenesis. Furthermore, inter-individual variability in genetic background and other confounding factors may obscure C9-ALS–specific chromatin disorganization. To directly assess whether the C9-STR expansion is the primary driver of C9-ALS gained loops, we controlled for genetic background by differentiating isogenic hiPSC lines with and without the C9-STR expansion^24–26^ into neurons, and profiled their 3D chromatin architecture using Micro-C^27^ (**Figure 2A**, N=6). We validated the presence of a G4C2 repeat expansion of ∼3kb (i.e., ∼500 repeats) in C9-STR hiPSCs, while no such band was detected in the isogenic control line (**Figure S4A**).

**Figure 2.**
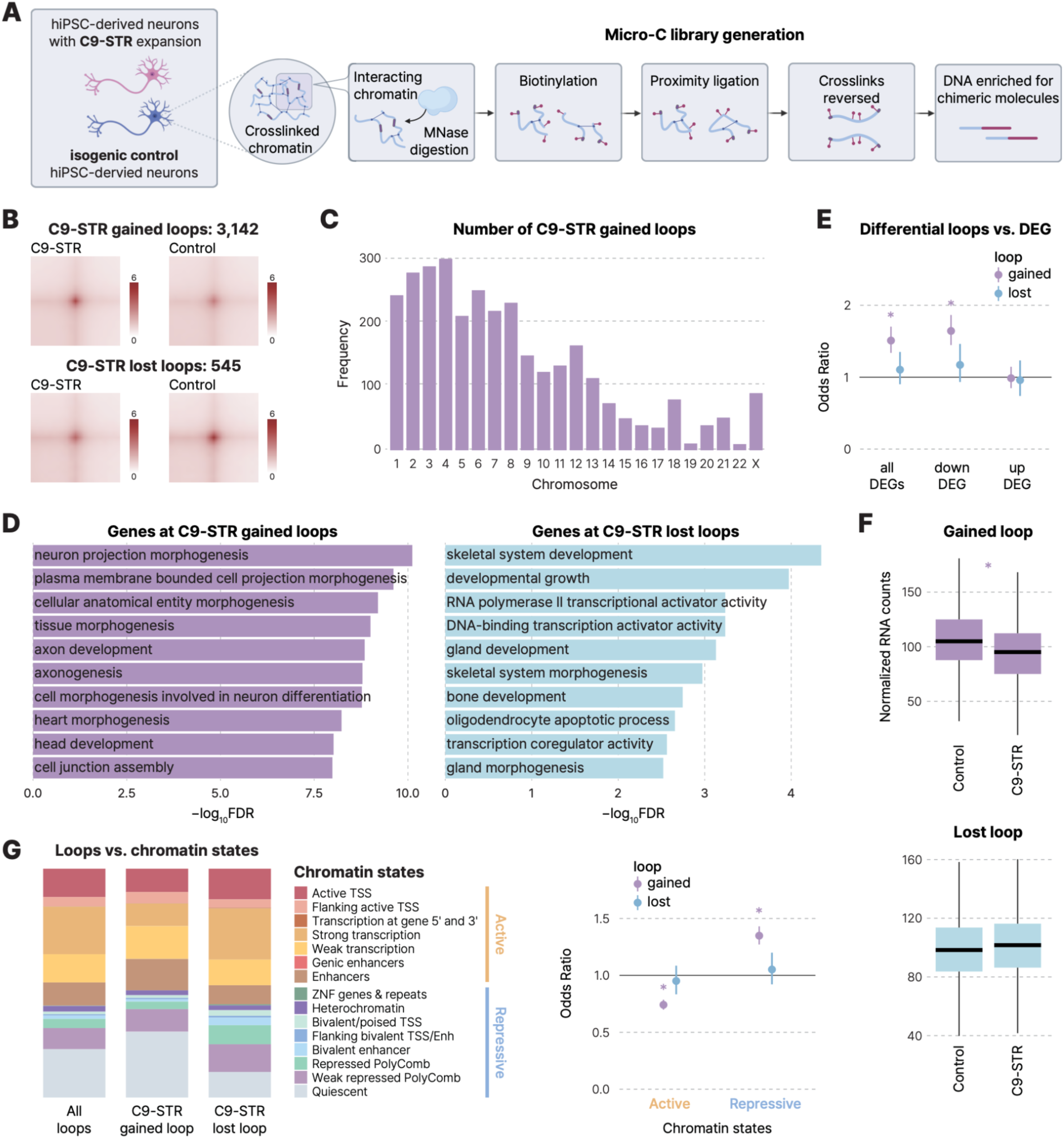
C9-STR expansion is associated with ectopic chromatin loops that are preferentially anchored at genes downregulated in C9-STR neurons. **A.** Genome-wide chromosome conformation maps were generated from hiPSC-derived neurons with the C9-STR expansion and isogenic controls. **B.** APA of C9-STR gained and lost loops. **C.** Genomic distribution of C9-STR gained loops, showing widespread distribution across chromosomes. **D.** Gene ontology analysis of genes anchored at C9-STR gained loops (left) and lost loops (right). **E.** C9-STR gained loops are enriched for downregulated genes in C9-STR neurons. Two-sided Fisher’s exact test, *FDR<0.05. **F.** Genes anchored at C9-STR gained loops were significantly downregulated in C9-STR neurons compared to isogenic controls, whereas genes anchored at C9-STR lost loops did not show C9-STR-specific changes in expression. One-sided Wilcoxon rank sum test, *p<2.2×10^-16^ for gained loops, p=0.94 for lost loops. **G.** Loop anchors of C9-STR gained loops are enriched for repressive chromatin states relative to all loops and C9-STR lost loops. Two-sided Fisher’s exact test, *FDR<0.05.

hiPSC-derived neurons exhibited a chromatin architecture similar to that of neurons isolated from the postmortem cortex (**Figure S4B**). Consistent with observations in postmortem neurons (**Figure 1B**), we did not observe disruption of TADs at the *C9orf72* locus in hiPSC-derived neurons carrying C9-STR expansion (hereafter referred to as C9-STR neurons to distinguish from C9-ALS postmortem neurons), compared to their isogenic counterparts lacking the pathogenic expansion (isogenic control neurons; **Figure S4C**). Similarly, interchromosomal interactions were comparable between C9-STR neurons and isogenic controls (**Figure S4D**).

Since C9-ALS neurons exhibited increased B-to-A compartment switching (**Figure S2B**), we next evaluated compartment changes in C9-STR neurons compared to isogenic controls. Overall, ∼17% of the genome exhibited A/B compartment switching (**Figure S5A**), a higher proportion than observed in postmortem neurons. This increase may reflect reduced technical variability in the isogenic hiPSC-derived models. Among regions with significant compartment changes, 39.96% transitioned from compartment B to A, while 33.95% transitioned from A to B, indicating only a minor shift toward more transcriptionally active compartments in C9-STR neurons (**Figure S5B**).

Next, we performed differential loop analysis between C9-STR and isogenic control neurons, identifying C9-STR gained loops (loops strengthened in C9-STR neurons) and C9-STR lost loops (loops weakened in C9-STR neurons). We observed a substantial imbalance, with 3,142 gained loops and only 545 lost loops in C9-STR neurons (**Figure 2B**). Both gained and lost loops were distributed across the genome (**Figure 2C**). In line with the marked increase in gained loops—and consistent with findings from postmortem neurons—C9-STR neurons exhibited more frequent TAD splitting than merging events (**Figure S5C**). These findings suggest that the C9-STR expansion drives widespread chromatin reorganization early in disease progression, rather than these changes representing secondary consequences of later-stage pathogenesis. Furthermore, C9-STR gained loops were anchored to genes enriched for neuronal differentiation and axonogenesis, whereas lost loops were linked to genes involved in transcriptional regulation, organ development, and oligodendrocyte differentiation (**Figure 2D**).

Because the excess of gained loops was observed in both C9-ALS neurons and hiPSC-derived C9-STR neurons, we next surveyed whether the increased TAD splitting detected in C9-ALS neurons was also recapitulated in the C9-STR neurons. Consistent with the postmortem findings, C9-STR neurons exhibited a higher frequency of TAD splitting compared to TAD merging events (**Figure S5C**). These splitting events significantly overlapped with C9-STR gained loops, suggesting that ectopic loop formation may be associated with local domain reorganization (**Figure S5C**).

### Relationship between C9-STR driven 3D genome disorganization and transcriptomic alterations

To investigate the relationship between C9-STR expansion-mediated chromatin disorganization and transcriptional regulation, we generated RNA-seq libraries from C9-STR and isogenic control neurons. Unlike in postmortem neurons (**Figure S2E**), regions involved in compartment switching were enriched for differentially expressed genes (DEGs; **Figure S5D, Table S4**). However, downregulated genes in C9-STR neurons were enriched in regions that underwent both A-to-B and B-to-A compartment switching, which contradicts the expectation that B-to-A transitions correspond to increased transcriptional activity. Therefore, we conclude that C9-STR–mediated compartment changes are not strongly linked to the molecular neuropathology of C9-ALS.

In contrast, C9-STR expansion-associated chromatin looping was closely linked to transcriptomic alterations. Genes anchored at C9-STR gained loops were significantly enriched for those downregulated in C9-STR neurons (**Figure 2E**), consistent with observations in postmortem neurons (**Figure 1G**). Moreover, these genes exhibited reduced expression relative to isogenic controls (**Figure 2F**). In comparison, C9-STR lost loops did not display enrichment for DEGs or notable changes in transcript levels (**Figure 2E-F**).

Although loop gains are often associated with increased transcriptional activity^19^, our data from both postmortem and hiPSC-derived neurons indicate the opposite: C9-STR gained loops were associated with transcriptional repression (**Figures 1G, 2E-F**). To explore this discrepancy, we examined the chromatin states of all loops, C9-STR gained loops, and C9-STR lost loops. Compared to all loops, C9-STR gained loops were more frequently anchored to repressive, particularly quiescent, chromatin regions. They were also less likely to be associated with active chromatin states, especially those linked to strong transcription (**Figure 2G**). In contrast, C9-STR lost loops were neither enriched nor depleted for active or repressive chromatin states (**Figure 2G**).

Together, the enrichment of downregulated genes at C9-STR gained loop anchors, observed consistently in both postmortem and hiPSC-derived neurons, highlights ectopic loop formation as a potential mechanistic driver of transcriptional dysfunction in C9-ALS^13–15^. C9-STRs may promote the formation of aberrant chromatin loops that juxtapose genomic regions not typically engaged in regulatory interactions, potentially leading to transcriptional misregulation. Notably, C9-STR gained loops were preferentially anchored at genes associated with neuronal function (**Figure 2D**), suggesting that their aberrant engagement may contribute to C9-ALS molecular pathology by disrupting neuronal gene expression and function.

### Transcribed C9-STRs interact with DNA

We next explored the mechanism by which the C9-STR expansion induces widespread chromatin remodeling. Since ectopic loops were distributed across the genome rather than restricted to the *C9orf72* locus (**Figures 1E, 2C**), we reasoned that this global chromatin rewiring must involve diffusible factors capable of acting throughout the nucleus. We therefore hypothesized that C9-STR RNAs, transcribed from the *C9orf72* locus, could disperse across the nuclear space and engage in molecular interactions that influence chromatin structure.

Growing evidence suggests that RNA-DNA interactions can orchestrate higher-order chromatin organization^28–31^. However, the contribution of such interactions in C9-ALS remains largely unexplored. To study whether C9-STR RNAs have a potential to bind to DNA, we performed capture hybridization analysis of RNA targets (CHART)^32,33^, a technique that isolates endogenous RNAs of interest along with their associated chromatin, enabling genome-wide mapping of RNA-DNA interactions.

We designed biotinylated antisense oligonucleotides targeting C9-STR RNAs (CCCCGG_4_ or C4G2 probes, see **Methods**) to isolate potential DNA interaction partners via CHART (**Figure 3A**). Because these C4G2 probes may also hybridize to random GC-rich sequences, we performed CHART in two conditions: (1) in C9-STR neurons, where the probes are expected to specifically target C9-STR RNAs, and (2) in isogenic control neurons, where the probes would primarily bind off-target GC-rich regions. As expected, we observed elevated CHART signals at the *C9orf72* locus in C9-STR neurons relative to isogenic controls, reflecting the presence of the STR expansion (**Figure 3B**).

**Figure 3.**
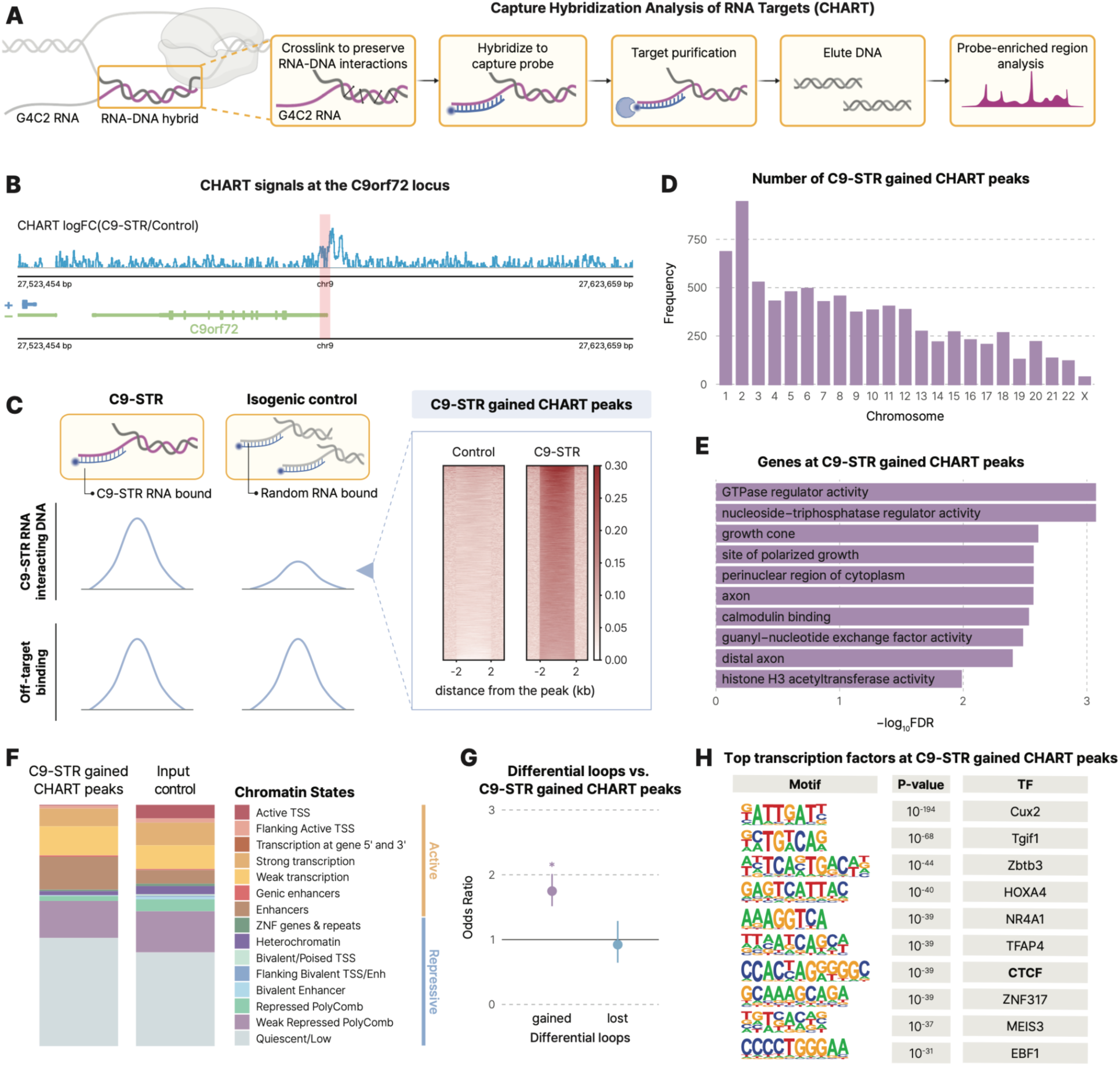
C9-STR RNAs preferentially bind genomic regions enriched for C9-STR gained loops. **A.** CHART was performed in hiPSC-derived neurons with the C9-STR expansion and isogenic controls to capture C9-RNA genomic targets. **B.** Log fold-change (logFC) of CHART signals at the *C9orf72* locus in C9-STR neurons compared to isogenic controls. **C.** Definition of C9-STR gained CHART peaks (left) and their enrichment in C9-STR neurons compared to controls (right). **D.** Genome-wide distribution of C9-STR gained CHART peaks, showing widespread distribution across chromosomes. **E.** Gene ontology enrichment analysis of genes located at C9-STR gained CHART peaks. **F.** Comparison of chromatin states between C9-STR gained peaks and input control peaks (no probe). C9-STR gained CHART peaks show an increased proportion of quiescent chromatin relative to the input control. **G.** C9-STR gained loops are significantly associated with C9-STR gained CHART peaks. Two-sided Fisher’s exact test, *FDR<0.05. **H.** Motif enrichment analysis identifying transcription factors enriched at C9-STR gained CHART peaks.

To identify genome-wide C9-STR RNA-associated RNA-DNA interactions, we focused on regions that show at least a 2-fold increase in CHART signal in C9-STR neurons relative to isogenic controls, consistently across two biological replicates (**Figure 3C, Table S5**). We refer to these regions as C9-STR gained CHART peaks. In total, we detected 8,059 gained CHART peaks (**Figure 3C**). Similar to C9-STR gained loops, gained CHART peaks were distributed throughout the genome (**Figure 3D**), supporting our hypothesis that C9-STR RNAs can disperse across the nucleus and seed DNA interactions at ectopic loci.

We next characterized potential biological functions of C9-STR gained CHART peaks. Gained CHART peaks were enriched at the promoters of genes involved in axonal growth (**Figure 3E**), mirroring the gene ontology terms enriched for C9-STR gained loops (**Figure 2D**). Furthermore, similar to the gained loops (**Figure 2G**), C9-STR gained CHART peaks more frequently overlapped with quiescent chromatin regions compared to input controls (**Figure 3F, Figure S6A**), suggesting that C9-STR RNAs preferentially bind genomic loci not previously engaged in regulatory interactions. Together, these results support the idea that C9-STR RNAs may associate with genomic regions critical for neuronal function and engage them in ectopic regulatory interactions, ultimately leading to transcriptomic dysregulation (**Figure 2E-F**).

Intrigued by these findings, we next asked whether C9-STR gained CHART peaks are enriched at C9-STR gained loops compared to stable loops. Indeed, gained CHART peaks were significantly enriched at the anchors of C9-STR gained loops, but not at the anchors of C9-STR lost loops (**Figure 3G**). To investigate how C9-STR RNA-mediated DNA interactions might contribute to chromatin loop formation, we performed transcription factor (TF) motif enrichment analyses using HOMER (**Table S6**)^34^. Notably, one of the top TF motifs enriched at gained CHART peaks was for CTCF, a well-known loop-inducing factor (**Figure 3H**), providing a potential mechanistic link between C9-STR RNAs and chromatin loop formation.

### C9-STR RNAs facilitate CTCF binding and ectopic loop formation

Given the unexpected enrichment of CTCF motifs at C9-STR gained CHART peaks, we performed CTCF CUT&RUN in C9-STR neurons and their isogenic controls to determine whether C9-STR RNA-bound genomic regions are also capable of recruiting CTCF (**Figure S6B**). In addition to motif enrichment (**Figure 3G**), C9-STR gained CHART peaks were significantly enriched for CTCF peaks (**Figure 4A**), suggesting that C9-STR RNAs may facilitate CTCF recruitment to ectopic genomic sites.

**Figure 4.**
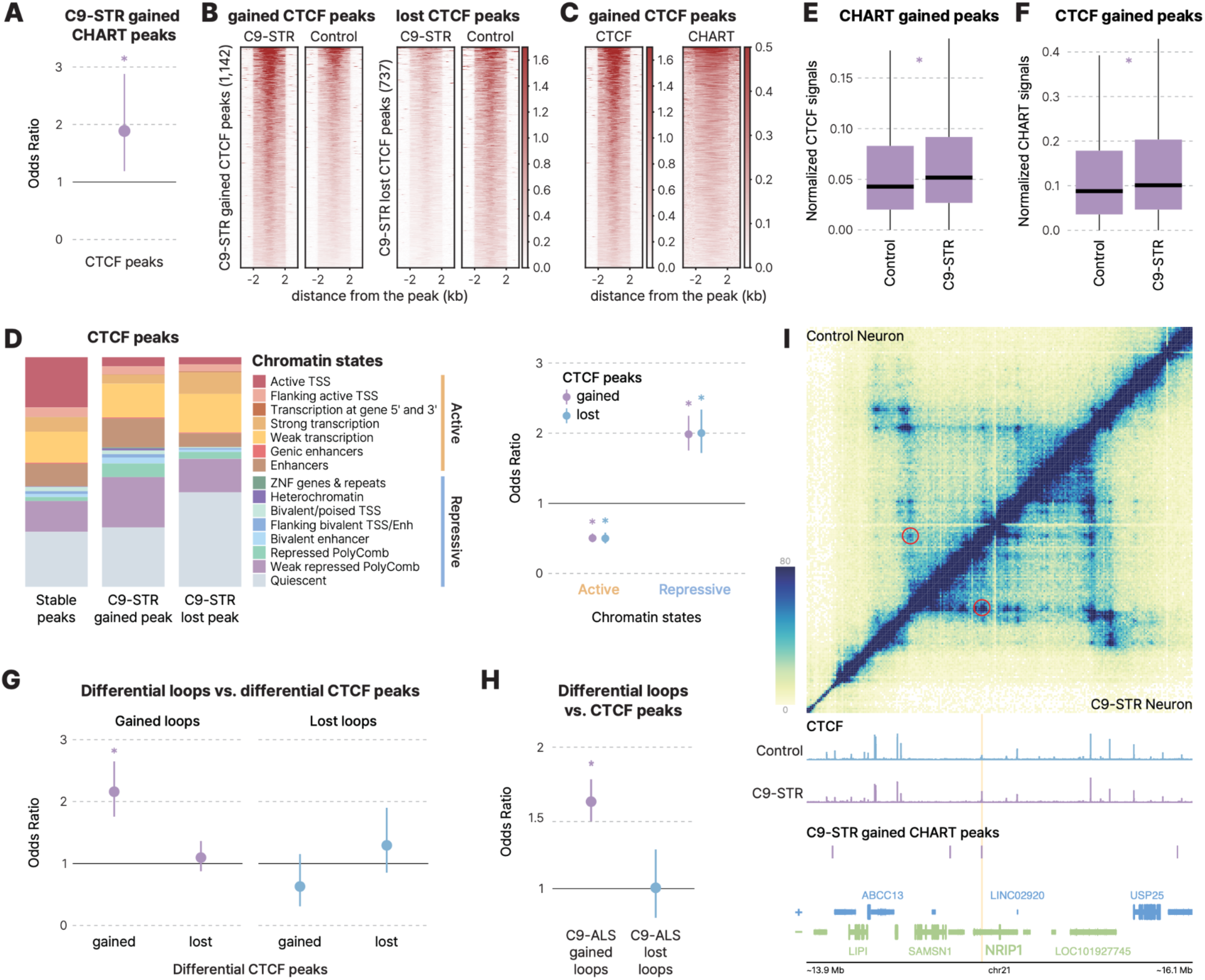
Relationship between CHART, CTCF, and loops in C9-STR neurons. **A.** C9-STR gained CHART peaks are enriched for CTCF peaks. Two-sided Fisher’s exact test, *p=0.0052. **B.** Gained and lost CTCF peaks in C9-STR neurons. **C.** CHART signals are enriched at C9-STR gained CTCF peaks in C9-STR neurons. **D.** C9-STR gained and lost CTCF peaks are more frequently associated with repressive chromatin states and depleted from active states relative to stable CTCF peaks. Two-sided Fisher’s exact test, *FDR<0.05. **E.** C9-STR gained CHART peaks show stronger CTCF signals in C9-STR neurons compared to isogenic controls. One-sided Wilcoxon rank sum test, *p<2.2×10^-16^. **F.** C9-STR gained CTCF peaks exhibit stronger CHART signals in C9-STR neurons compared to isogenic controls. One-sided Wilcoxon rank sum test,*p=1.95×10^-5^. **G.** C9-STR gained loops, but not lost loops, are enriched for C9-STR gained CTCF peaks. Two-sided Fisher’s exact test, *FDR<0.05. **H.** C9-ALS gained loops are enriched for CTCF peaks compared to stable loops. Two-sided Fisher’s exact test, *FDR<0.05. **I.** A C9-STR gained loop (red circle) colocalizes with a C9-STR gained CTCF peak and CHART peak (orange bar) at the *NRIP1* locus. Chromatin organization in isogenic control neurons is shown in the top left, while chromatin organization in C9-STR neurons is shown in the bottom right.

We next asked whether overall CTCF occupancy differs between C9-STR neurons and isogenic controls. Although the total number of CTCF peaks was lower in C9-STR neurons, differential analysis based on CTCF signal intensity identified 1,142 C9-STR gained CTCF peaks (i.e., peaks significantly strengthened in C9-STR neurons) and 737 C9-STR lost CTCF peaks (**Figure S6C, Table S7**). These results indicate that, despite a global reduction in peak count, many genomic loci exhibit increased CTCF binding in C9-STR neurons. This redistribution of CTCF binding is consistent with the observed increase in C9-STR gained loops (**Figure 4B, S6C**). Both gained and lost CTCF peaks were broadly distributed across the genome, with no specific enrichment on chromosome 9, where the *C9orf72* locus resides (**Figure S6D**).

Consistent with the enrichment of CTCF at C9-STR gained CHART peaks (**Figure 4A**), we observed that C9-STR gained CTCF peaks were reciprocally enriched for CHART signals in C9-STR neurons (**Figure 4C**). Together, these results support a model in which C9-STR RNAs act as molecular scaffolds that promote ectopic CTCF binding and contribute to chromatin misfolding in C9-ALS.

CTCF peaks were predominantly enriched at the promoters of genes implicated in synaptic function, consistent with its well-established role in neuronal gene regulation (**Figure S6E**)^35,36^. In contrast, C9-STR gained CTCF peaks showed no enrichment for gene ontology terms, suggesting that C9-STR RNAs may recruit CTCF to ectopic genomic regions not typically engaged in regulatory activity. Supporting this hypothesis, these gained CTCF peaks were more frequently associated with repressive chromatin states and less frequently with active states (**Figure 4D**).

Interestingly, C9-STR lost CTCF peaks were enriched at promoters of genes involved in neuronal and synaptic function. This suggests that CTCF, which would normally bind to these sites to regulate neuronal gene expression, may be displaced and aberrantly recruited to ectopic regions in the presence of C9-STR RNA (**Figure S6E**). Similar to C9-STR gained CTCF peaks, lost CTCF peaks were also more frequently associated with repressive chromatin states and less frequently with active states compared to all CTCF peaks (**Figure 4D**), indicating that altered CTCF occupancy in C9-STR neurons predominantly involves repressive chromatin regions.

Prompted by these findings, we explored whether C9-STR RNA-bound loci are susceptible to altered CTCF binding. Measuring CTCF occupancy at C9-STR gained CHART peaks revealed significantly higher CTCF signals in C9-STR neurons compared to isogenic controls (**Figure 4E**). Conversely, when we measured CHART signals at C9-STR gained CTCF peaks, we also observed increased signals in C9-STR neurons relative to controls (**Figure 4F**). Together, these results suggest a reciprocal relationship in which C9-STR RNA-bound loci are more likely to recruit CTCF, thereby linking RNA-DNA interactions to ectopic CTCF binding.

To assess the functional consequences of altered CTCF binding, we next examined its relationship to 3D genome organization. C9-STR gained CTCF peaks were significantly enriched at the anchors of C9-STR gained loops, but not at the anchors of C9-STR lost loops (**Figure 4G**). In contrast, C9-STR lost CTCF peaks were not enriched at the anchors of either C9-STR gained or lost loops (**Figure 4G**).

We extended this analysis to C9-ALS neurons by intersecting differential chromatin loops with CTCF ChIP-seq data from human neurons^37^. Given the well established CTCF enrichment at loop anchors, we compared differential loops to stable loops, which served as a baseline for CTCF occupancy. CTCF peaks were more frequently observed at C9-ALS gained loops compared to stable loops (**Figure 4H**). In contrast, C9-ALS lost loops did not exhibit significant differences in CTCF occupancy relative to stable loops (**Figure 4H**). Together, these findings suggest that CTCF is preferentially associated with gained loops—both in C9-STR and C9-ALS—and may act as a key mediator of ectopic loop formation in C9-ALS.

One illustrative example of colocalization between C9-STR gained loops, CHART peaks, and CTCF peaks occurs at the *NRIP1* locus (**Figure 4I**), where all three features converge. *NRIP1* has been implicated in neuromuscular phenotypes: knockout mice exhibit muscular abnormalities and reduced regenerative capacity^38^, and human loss-of-function variants have been associated with neurodevelopmental disorders, including spastic cerebral palsy ^39^. This example underscores a potential link between STR-driven chromatin misfolding and transcriptional dysregulation of a gene implicated in motor system vulnerability.

## Discussion

Our study proposes a previously unrecognized mechanism underlying ALS pathogenesis, whereby the C9-STR expansion induces widespread reorganization of the 3D genome, characterized by a pronounced increase in chromatin loops, particularly in neurons.

The human genome harbors numerous GC-rich, repetitive regions that are prone to forming G4 structures when single-stranded during replication or transcription. When one DNA strand folds into a G4 structure, the complementary strand becomes more accessible and can hybridize with RNA, promoting the formation of R-loops, three-stranded nucleic acid structures formed when RNA hybridizes to its template DNA strand^40^. Once established, G4 structures and R-loops can mutually stabilize each other by preventing reannealing, creating highly persistent RNA-DNA structures^41^. Importantly, both R-loops and G4 structures have been shown to recruit CTCF, a loop-inducing TF central to 3D chromatin architecture^7,42,43^. Given these features of GC-rich DNA and our current findings, we propose that transcribed C9-STRs act as diffusible scaffolds that engage complementary GC-rich genomic regions, establishing RNA-DNA interactions that predispose these loci to structural reorganization.

We demonstrated that C9-STR RNAs engage in widespread RNA-DNA interactions across the genome, and these RNA-bound regions exhibit reciprocal enrichment with CTCF binding. We therefore propose that C9-STR RNA-mediated recruitment of CTCF provides a mechanistic basis for the widespread gain of chromatin loops in C9-ALS neurons^8^ (**Figure 5**). These gained loops are preferentially anchored in repressive chromatin states or establish new regulatory linkages with previously unengaged genomic regions, thereby contributing to the widespread transcriptional downregulation observed in C9-ALS brains^13–15^.

**Figure 5.**
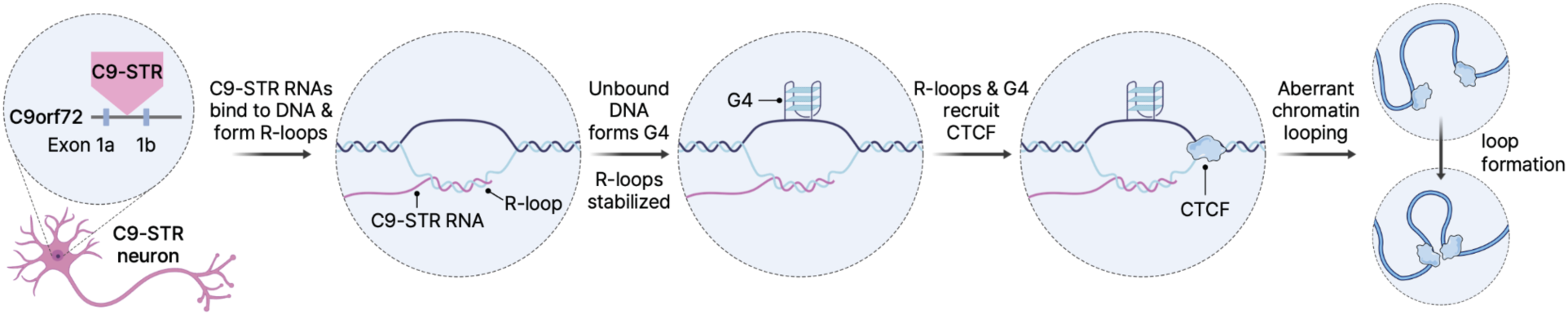
Hypothetical model of how C9-STR RNAs induce ectopic chromatin loops. Expanded C9-STRs are transcribed into RNA, which can hybridize with complementary DNA sequences to form RNA-DNA hybrids, or R-loops. The displaced single-stranded DNA strand is prone to folding into G4 structures. Together, these R-loops and G4 structures can recruit loop-inducing TFs such as CTCF, promoting the formation of ectopic chromatin loops in C9-STR neurons.

Recently, R-loops have been proposed as therapeutic targets for STR-related disorders such as FXS^44^. Our findings provide fundamental insight into this strategy by demonstrating that RNA-mediated DNA interactions can have broader implications for chromatin topology and gene regulation, potentially serving as key intervention points for STR-driven diseases like C9-ALS. Mapping high-resolution R-loop landscapes in C9-ALS neurons using orthogonal approaches such as MapR or DRIP-seq will be essential to validate the involvement of persistent DNA-RNA hybrid structures in disease etiology.

Moreover, it remains unknown whether C9-STR RNAs bind DNA stochastically or are guided by specific sequence or structural cues. Elucidating the regulatory logic and specificity of C9-STR RNA-DNA interactions could uncover fundamental mechanisms of RNA-guided genome organization and reveal new dimensions of C9-ALS pathophysiology. Finally, as the role of STR expansions in ALS becomes increasingly recognized^45^, investigating whether transcribed STRs broadly impact various aspects of 3D genome architecture represents an important direction for future research.

Overall, our findings demonstrate that C9-STR expansions reshape neuronal chromatin architecture through RNA-guided mechanisms, directly connecting repeat-driven genome misfolding to the transcriptomic alterations underlying C9-ALS.

## Methods

### Generation of Hi-C libraries from postmortem brain cells

Nuclei were isolated from the visual cortex of three individuals with C9-ALS and three neurotypical controls. Neuronal (NeuN-positive) and glial (NeuN-negative) nuclei were separated using fluorescence-activated nuclei sorting (FANS), as previously described^16^. For each sample, ∼1 million nuclei were processed to generate Hi-C libraries following established protocols^16,17,46^. Briefly, sorted nuclei were fixed in 1% formaldehyde to preserve chromatin interactions. Crosslinked chromatin was then digested with MboI (NEB, R0147S), followed by end repair, biotin labeling, and proximity-based ligation. The resulting DNA was sheared to 300-600 base-pair (bp) fragments, and biotinylated fragments were selectively captured via streptavidin pulldown. Illumina sequencing adapters were ligated to the recovered fragments, which were then amplified by PCR. Hi-C libraries were sequenced using 100 bp paired-end reads on an Illumina HiSeq.

### Generation of hiPSC-derived neurons with and without the C9-STR expansion

We obtained hiPSC-derived neurons harboring the C9-STR expansion (CS52iALS-C9nxx) and their isogenic control (CS52iALS-C9n6.ISOxx) from the Cedars Sinai hiPSC Core Facility. hiPSC lines were maintained on Matrigel-coated dishes (Corning, 354480) in mTeSR Plus Medium (Stem Cell Technologies) and passaged every 3-4 days with 0.5 mM EDTA, as previously described ^47^.

Cortical neuronal differentiation was performed following a previously described protocol^48^. Cells were cultured through the neural progenitor cell (NPC) stage until day 12, then expanded for 7 days in Neuronal Expansion Medium (1:1 Advanced DMEM/F-12, 12634028; Neural Induction Supplement, A1647801) supplemented with 20 ng/mL FGF (PeproTech, 100-18B). NPCs were further expanded in a bioreactor until the desired yield was reached.

For neuronal maturation, NPCs were plated onto poly-D-lysine/laminin-coated plates and maintained in Cortical Neuron Maturation Medium consisting of 1:1 Advanced DMEM/F-12 and Neurobasal, supplemented with 1x GlutaMAX, 100 mM β-mercaptoethanol, 1x B27, 0.5x N2, 1x non-essential amino acids (NEAA), and 2.5 µg/mL insulin (Sigma-Aldrich, I9278). Maturation factors included 10 ng/mL BDNF, 10 ng/mL GDNF, and 10 µM DAPT. Cells were cultured under these conditions for 3-4 weeks prior to downstream genomic assays. Neuronal identity was confirmed by expression of NPC markers (PAX6, SOX2, SOX1) and mature cortical neuronal markers (MAP2, SATB2).

For cryopreservation, matured neurons were dissociated with TrypLE supplemented with 25 mM trehalose and 50 µg/mL DNase. After 7 minutes, cells were collected and TrypLE reaction was quenched with soybean trypsin inhibitor. Cells were pelleted by centrifugation, resuspended in fresh media, filtered, counted, and aliquoted for freezing. The cryopreservation medium contained 25 mM trehalose and 10% DMSO. Aliquots (1 mL per cryovial) were transferred to Mr. Frosty containers and frozen at −80 °C for 24 hours before storage in the vapor phase of liquid nitrogen until use. Unless otherwise specified, reagents were obtained from Thermo Fisher Scientific.

### Repeat-PCR to assess G4C2 repeat length in hiPSCs

Repeat-PCR was used to confirm the length of the G4C2 repeat expansion in C9-STR hiPSCs.

Genomic DNA was isolated from hiPSCs using the Zymo Quick-DNA Miniprep Plus Kit (D4068). Isolated DNA was digested with HindIII in a reaction buffer containing 50 mM Tris-Cl (pH 8.8), 1.5 mM MgCl₂, 22 mM (NH₄)₂SO₄, and 0.2% Triton X. The digested DNA served as the template for a Repeat-PCR reaction, prepared with the following components: 50 mM Tris-Cl (pH 8.8), 1.5 mM MgCl₂, 22 mM (NH₄)₂SO₄, 0.2% Triton X, 3.3 M betaine, 2.67% DMSO, 0.45 mM dNTPs, 0.67 µM of each primer, and 0.02 U/µl Q5 DNA polymerase (NEB, M0491S)^49^.

PCR amplification was performed as follows: an initial denaturation at 98 °C for 5 minutes, followed by 7 touchdown cycles of 97 °C for 35 seconds, 72-62 °C for 35 seconds (–1.6 °C per cycle), and 82 °C for 3 minutes. This was followed by 30 cycles of 97 °C for 35 seconds, 56 °C for 15 seconds, 64 °C for 20 seconds, and 82 °C for 2 seconds, interspersed with 18 minicylces consisting of 84 °C for 2 seconds (30% ramp rate) and 82 °C for 2 seconds (100% ramp rate). A final extension step was carried out at 72 °C for 10 minutes, and reactions were held at 4 °C^50^. PCR products were resolved on agarose gels and visualized by ethidium bromide staining.

### Generation of Micro-C libraries from hiPSC-derived neurons

Micro-C was performed using the Dovetail Genomics Cantata Bio Micro-C kit (Dovetail Proximity Ligation Core Box 1, PN DG-REF-001; Dovetail Micro-C Module Box 2, PN DG-NUC-001; Dovetail Library Module, PN DG-LIB-001), following the manufacturer’s user guide (version 1.0). Isogenic hiPSC-derived neurons, either harboring or lacking the C9-STR expansion, were used as input. Six biological replicates were prepared for each condition, using 1.5 million cells per replicate.

Cells were crosslinked using double fixation with 3 mM DSG followed by 1% formaldehyde, preserving native chromatin architecture. Crosslinked chromatin was digested using 1.5 μL MNase at 22°C for 25 minutes. The reaction was stopped by adding 0.5 M EGTA, and cell lysis was performed using 20% SDS to release chromatin contents. To verify digestion efficiency, an aliquot of the lysate was purified using the Zymo DNA Clean and Concentrator-5 Kit (D4004). According to the Dovetail protocol, samples showing 40-70% mononucleosome content were deemed successfully digested and advanced to proximity ligation. Next, chromatin capture Beads were used to isolate chromatin fragments. End-repair was performed, followed by ligation to biotinylated bridge adapters. Proximity ligation was then carried out to link spatially adjacent chromatin fragments. After reversing the crosslinks, DNA was purified with SPRIselect beads, and Illumina paired-end sequencing adapters (xGen UDI Primers Plate 1, 8nt, #10005922) were ligated. Biotinylated ligation products were enriched using streptavidin beads, followed by library amplification via PCR. Final libraries were size-selected using SPRIselect beads to enrich for fragments in the 350-1,000 bp range. Sequencing was performed on the Illumina NovaSeq 6000 platform, generating approximately 4 billion 150 bp paired-end reads.

### Generation of RNA-seq libraries from hiPSC-derived neurons

RNA was extracted from 1 million hiPSC-derived neurons per sample for each of 6 biological replicates per condition (C9-STR neurons and isogenic control neurons, n=12 samples total) using the Zymo Quick-RNA Miniprep Kit (R10564). RNA concentration was measured using the Qubit RNA High Sensitivity Assay Kit (Invitrogen, Q32852), and RNA quality was assessed using the RNA Integrity Number (RIN) obtained via the Agilent High Sensitivity RNA ScreenTape Assay (5067-5579). Library preparation was performed by the UNC Integrated Genomics Core High-Throughput Sequencing Facility (RRID: SCR_022620) using the KAPA Stranded mRNA-Seq Kit (Roche Diagnostics, 07962193001), following the manufacturer’s protocol. Prepared libraries were sequenced on the Illumina NovaSeq X 1.5B platform, generating 100 bp paired-end reads at an average depth of 50 million read pairs per sample.

### CHART in hiPSC-derived neurons

CHART was performed on isogenic hiPSC-derived neurons with and without the C9-STR expansion (two biological replicates per condition; 30-35 million cells per replicate). Cells were first fixed with 1% PFA for 10 minutes at room temperature to preserve chromatin architecture, followed by nuclei isolation. A second crosslinking step was performed on isolated nuclei using 3% formaldehyde for 10 minutes at room temperature. After washing, nuclei were resuspended in sonication buffer (50 mM HEPES-OH pH 7.5, 75 mM NaCl, 0.5% N-lauroylsarcosine, 0.1% sodium deoxycholate, 0.1 mM EGTA) and sonicated using a Covaris M220 Focused-ultrasonicator (75 Peak Incident Power, 10% Duty Factor, 200 Cycles per Burst, 3 minutes) to obtain 2-10 kb chromatin fragments. Crosslinks were then reversed, and the DNA was purified and assessed for fragmentation. Chromatin extracts were incubated with Dynabeads MyOne Streptavidin C1 beads (50 μL beads per 750 μL chromatin extract) for 1 hour at room temperature to minimize non-specific binding prior to hybridization.

For capture hybridization, the extracts were incubated overnight at room temperature with 250 pmol of a custom biotinylated DNA probe targeting the G4C2 motif (5’ - CCC CGG CCC CGG CCC CGG CCC CGG /Sp18//3BioTEG/-3’; 24 bp, 100 nmol, IDT). An input control (no probe added) was included for each replicate. Following hybridization, the samples were incubated with 200 μL of Dynabeads MyOne Streptavidin C1 beads per 250 pmol of probe to capture the RNA-DNA complexes. After elution, both RNA and DNA were purified. DNA was then re-sonicated to a final size of 100-500 bp (50 Peak Incident Power, 10% Duty Factor, 200 Cycles per Burst, 70 seconds) using the Covaris M220. DNA libraries were prepared using the CUTANA CUT&RUN Library Prep Kit (EpiCypher, 14-1001) and sequenced on the Illumina NextSeq 2000 XLEAP P2 platform, generating 150 bp paired-end reads.

### Generation of CUT&RUN libraries from hiPSC-derived neurons with and without the C9-STR expansion

CUT&RUN was performed using the CUTANA CUT&RUN Kit (Epicypher, 14-1048) using 4 replicates per condition (C9-STR neurons and isogenic control neurons, totaling 8 samples). Briefly, nuclei were isolated from hiPSC-derived neurons following the nuclei isolation protocol outlined in the CUTANA CUT&RUN Kit. 550,000 nuclei per reaction were aliquoted and incubated with pre-activated concanavalin A-coated beads at room temperature for 10 min, followed by overnight antibody incubation in a cell permeabilization buffer containing 0.01% digitonin at 4°C. The following antibodies were used: rabbit IgG for negative control (Epicypher, 13-0042) and rabbit anti-CTCF (Epicypher, 13-2014). Following incubation, nuclei bound to concanavalin A-coated beads were permeabilized with a buffer containing 0.01% digitonin and incubated with the pAG-MNase fusion protein at room temperature for 10 min. After washing, chromatin-bound pAG-MNase cleavage was induced by addition of calcium chloride at a final concentration of 2mM. This reaction was carried out at 4°C for 2 h and then stopped by addition of a stop buffer (containing fragmented genomic Escherichia coli DNA as spike-in). Following fragmented DNA purification, Illumina sequencing libraries were prepared from 5 ng of purified DNA (calculated with the Qubit dsDNA HS Assay Kit) using the CUTANA CUT&RUN Library Prep Kit (EpiCypher 14-1001) according to the manufacturer’s recommendations. Libraries were sequenced on the Illumina NextSeq 2000 platform, producing approximately 50 million 75 bp paired-end reads per sample.

### Hi-C and Micro-C analysis

Read quality of fastq files was first assessed using FastQC (v0.12.1). Sequencing reads were then aligned to the GRCh38 reference genome using bwa mem (v0.7.17). Duplicate reads were removed with Pairtools (v1.1.3)^51^. Resulting BAM files were converted to multi-resolution contact matrices using cooler (v0.10.4) for downstream analysis.

### Comparison of brain-derived Hi-C and Micro-C datasets

We used HiCRep (v1.12.2)^52^ to evaluate similarities among control and C9-ALS samples derived from postmortem neuronal and glial Hi-C libraries, as well as hiPSC-derived neuronal Micro-C libraries. The stratum-adjusted correlation coefficient (SCC) matrix was computed at 100 kb resolution using a smoothing parameter of *`h`*`=3`.

### Differential compartment analysis

We performed differential compartment analysis using dcHiC (v2.1)^11^, and called compartments at the resolution of 100kb. The `--cis` flag was used to identify A/B compartments from cis interaction matrices, and `--subnum 4` was applied to assign subcompartments. Transitions in compartment identity between control and C9-ALS/C9-STR samples were visualized using a Sankey diagram generated with the networkD3 R package (v0.4.1).

### Differential TAD analysis

TAD boundaries were identified using cooltools insulation (v0.7.1) at 40 kb resolution with a 160 kb window size. To detect changes in TAD architecture between conditions, differential TADs were identified using diffDomain^21^, which classifies TADs into six subtypes based on boundary dynamics. To evaluate the relationship between TAD reorganization and chromatin looping, we performed a two-sided Fisher’s exact test in R (v4.4.0) to assess the statistical significance of the overlap between TAD-splitting events and C9-STR gained loops in hiPSC-derived neurons.

### Differential loop analysis

Chromatin loops were identified genome-wide using Mustache (v1.0.1)^53^ at 10 kb resolution. For differential loop analysis, loop counts were extracted from cooler-formatted contact matrices and normalized by library size. A linear regression model was applied with biological condition as a covariate to assess condition-specific differences in loop strength. FDR correction was used to adjust *p*-values for multiple testing. Differential loops identified in both C9-ALS and C9-STR neurons were further validated using Aggregate Peak Analysis (APA), computed with the apa-analysis command from hicpeaks.

C9-ALS differential loops were compared against CTCF ChIP-seq data from neurons^37^. We used a two-sided Fisher’s exact test to assess whether gained or lost loops are enriched for overlap with CTCF peaks compared to stable loops.

### Cellular expression of *C9orf72*

To assess the cell type-specific expression profile of *C9orf72* across the human cortex, we accessed transcriptomic data from the UCSC Cell Browser’s integrated dataset, “A Meta-Atlas of the Developing Human Cortex,” which catalogs gene expression across diverse cortical cell populations.

### Differential gene expression analysis

RNA-seq reads were aligned to the reference genome using STAR (v2.7.11b)^54^, and gene-level quantification was performed using HTSeq (v2.0.5)^55^. Differential expression analysis was conducted using DESeq2 (v1.44.0)^56^, with a model that included replication and condition as design factors. A likelihood ratio test (LRT) was used to assess the effect of condition by comparing the full model (replication + condition) against a reduced model containing only replication. Using an adjusted p-value (p_adj_) < 0.05 and log2 fold change (LFC) > 0, we identified 4,828 upregulated DEGs. Similarly, 4,931 downregulated DEGs were identified using p_adj_ < 0.05 and LFC < 0.

We also obtained differentially expressed genes (DEGs) from postmortem neurons of individuals with C9-ALS using a previously published dataset^13^. We included the following neuronal subtypes: ALS Exc_deep, Exc_intermediate, Exc_superficial, Exc_unknown, Inh_ADARB2_Other, Inh_LAMP5, Inh_PVALB, Inh_SST, and Inh_VIP. Genes were classified as upregulated if they had p_adj_ < 0.05 and LFC > 0, and as downregulated if they had p_adj_ < 0.05 and LFC < 0 in any of the selected subtypes.

### Integration of DEGs with chromatin architectures

To compare DEGs with differential chromatin compartments, we first identified genes overlapping significant A-to-B or B-to-A compartment transitions. We then constructed a contingency table with the following categories: (i) genes overlapping both DEGs and compartment-transition regions, (ii) compartment-transition genes not overlapping with DEGs, (iii) DEGs not overlapping with compartment-transition regions, and (iv) genes in neither set. A two-sided Fisher’s exact test was performed to assess the statistical significance of the overlap. This analysis was repeated separately for upregulated and downregulated DEGs, and all p-values were adjusted using the FDR method.

To compare DEGs with differential TADs, we first identified genes overlapping either significant or stable TADs. A contingency table was constructed comparing the number of overlapping genes between DEGs and significant TADs versus those overlapping with stable TADs or neither. A two-sided Fisher’s exact test was performed to assess the enrichment of DEGs within significant TADs. This analysis was also conducted separately for upregulated and downregulated DEGs, and FDR-adjusted p-values were reported.

For the differential loop analysis, we identified genes overlapping gained, lost, or stable loops. Contingency tables were constructed to compare overlaps between DEGs and each loop category. Fisher’s exact tests were performed to assess the statistical significance of enrichment, and the analysis was repeated separately for upregulated and downregulated DEGs. All resulting p-values were FDR-corrected.

Finally, for the summation-based differential expression test, we identified all genes overlapping gained loops and quantified RNA-seq counts using `bedtools multicov`. Count matrices were normalized by samples, and the average expression for C9-STR and control samples was computed. A one-tailed Wilcoxon rank-sum test was used to determine whether gene expression was significantly higher in C9-STR samples compared to controls.

### CHART analysis

CHART reads were aligned to the GRCh38 reference genome using bwa mem (v0.7.17)^57^. Duplicate reads were removed with Picard (v3.1.1), and replicates were merged using Samtools (v1.6). Peaks were called using MACS2 (v2.2.9.1)^58^ with the following parameters: `-t STR_nodups_sorted.bam-f BAMPE-q 0.05-g hs --keep-dup all-B –SPMR`. Read counts per peak were quantified using bedtools multicov (v2.31.0) on the merged BAM files for each control and STR replicate. C9-STR gained CHART peaks were defined if the mean CPM-normalized read count in C9-STR samples was at least two-fold higher than that in control samples.

Motif enrichment analysis was conducted on the C9-STR gained CHART peaks using HOMER (v5.1) via the findMotifsGenome.pl tool^34^. We report both *de novo* and known motifs and defined motifs with p-values less than 1×10^-10^ as significant.

We defined the union of CHART input control peaks from control and C9-STR samples as the reference input set. Using the findOverlaps function from the IRanges R package (v2.38.1), we assessed the overlap between gained and stable loops with gained and input control CHART peaks. A two-sided Fisher’s exact test was performed to evaluate the statistical significance of enrichment between gained loops and gained CHART peaks. The same analysis was conducted for lost loops, and all p-values were adjusted for multiple testing using the FDR method.

### CUT&RUN analysis

CTCF CUT&RUN reads were preprocessed following the same pipeline as for CHART analysis. Replicates were merged separately for C9-STR and isogenic control conditions. Peaks were called using MACS2 (v2.2.9.1) with the following parameters: `-t CTCF_STR_isoCtrl.bam-c input_nodups_sorted.bam-f BAM-q 0.01-g hs --nomodel --shift 0 --extsize 200 --keep-dup all-B –SPMR`. Differential peaks were identified using DESeq2 (v1.44.0) with an LRT comparing a full model that adjusts for replication and condition against a reduced model that includes replication only. Log fold change shrinkage was performed using the apeglm method. C9-STR gained CTCF peaks were defined as those with p_adj_ < 0.05 and LFC > log₂(1.5), while C9-STR lost CTCF peaks were defined as p_adj_ < 0.05 and LFC < −log₂(1.5).

### Integrative analysis of CHART and CTCF profiles

To compare C9-STR gained CHART peaks with C9-STR gained CTCF peaks, we constructed a contingency table based on overlaps identified by the findOverlaps function from the IRanges R package (v2.38.1), categorizing peaks as gained or not-gained for both CHART and CTCF datasets. A two-sided Fisher’s exact test was performed to evaluate the statistical significance of the enrichment between C9-STR gained CHART and CTCF peaks.

For the summation-based differential signal analysis, we first calculated the mean CTCF signal in control and C9-STR samples at each CHART-gained peak region. A one-sided Wilcoxon rank-sum test was then used to evaluate whether the mean CTCF signal was significantly higher in STR samples compared to controls. Similarly, we calculated the mean CHART signal in control and C9-STR samples at each CTCF-gained peak region and applied a one-sided Wilcoxon rank-sum test to assess whether CHART signal was significantly increased in C9-STR samples compared to isogenic controls.

### Comparison of CTCF with chromatin loops

We used the findOverlaps function in the IRanges R package (v2.38.1) to assess overlap between gained vs. stable loops and gained vs. not-gained CTCF peaks. A two-sided Fisher’s exact test was used to test the statistical significance of the enrichment between gained loops and gained CTCF peaks. This analysis was repeated across all combinations of loop and CTCF peak status: gain-gain, gain-loss, loss-gain, and loss-loss.

### Genome segmentation analysis using ChromHMM

We used the ChromHMM core 15-state model from stem cell-derived neurons (E010) as a reference to annotate loop and peak regions^59^. For regions overlapping multiple chromatin states, the state with the lowest numerical label (i.e., the most active state) was assigned. The proportion of loop or peak regions falling within each chromatin state was then calculated.

To assess enrichment of loop/peak regions in active vs. repressive chromatin states, we performed Fisher’s exact tests. Active states included: active transcription start sites (TSS), flanking active TSS, transcription at gene 5′ and 3′ ends, strong transcription, weak transcription, genic enhancers, and enhancers. Repressive states included: ZNF genes/repeatsm heterochromatin, bivalent/poised TSS, flanking bivalent TSS/enhancers, bivalent enhancers, repressed Polycomb, weak repressed Polycomb, and quiescent.

### Gene Ontology (GO) enrichment analysis

GO enrichment analysis was conducted using gprofiler2 (v0.2.3), using a list of genes (GENCODE version 38) whose promoters were either anchored at chromatin loops or overlapped with CTCF/CHART peaks. All protein-coding genes were used as the background set. The top GO terms with FDR < 0.05 were reported.

### Data visualization

Hi-C and Micro-C contact matrices, as well as CTCF and CHART signals, were visualized using the plotgardener R package (v1.10.2)^60^.

We also used deepTools (v3.5.2) to compute signal profiles at gained or lost peak regions in the merged C9-STR samples and isogenic controls. We first ran `computeMatrix scale-regions` with the following parameters:-`-beforeRegionStartLength 2000`, `--regionBodyLength 5000`, `--afterRegionStartLength 2000`, `--sortRegions descend`, and `--sortUsing mean`, generating a matrix file sorted by mean signal intensity in descending order. Using the resulting sorted BED file, we then generated a matching matrix for the merged control samples with the same parameters. Heatmaps for the STR and control samples were plotted separately using `plotHeatmap`, and the final side-by-side display was created using the `convert +append` command.

## Acknowledgements

We thank members of the Won lab and Dr. Scott Behie from Keio University for helpful discussions and comments about this paper. This research was supported by the McKnight Foundation (H.W.), ALS Association (H.W.), VA Merit grant (BX005585, S.D.), NIA (R61AG094646, R01NS128523, H.W.), and NIGMS (R35GM153293, J.M.C.).

## Author Contributions

W.M., S.D., and H.W. conceived the idea of profiling 3D chromosome conformation in C9-ALS and designed the study. S.D. and A.K. sorted neurons and glia from postmortem brains. W.M. generated Hi-C libraries from sorted neurons and glia. A.A.B and A.B.M.G. maintained and differentiated hiPSCs with and without the C9-STR expansion into neurons under A.S.B.’s supervision. T.U.A. generated Micro-C libraries from hiPSC-derived neurons. L.A. generated CUT&RUN and RNA-seq libraries from hiPSC-derived neurons. T.U.A. created CHART libraries from hiPSC-neurons under J.M.C.’s supervision. S.U. performed repeat-PCR under H.G.L.’s supervision. J.W. and E.H. performed genomic analysis on Hi-C and Micro-C libraries. J.W. performed integrative analysis across Micro-C, CUT&RUN and CHART analysis. T.U.A., J.W., L.A. jointly generated figures. H.W., T.U.A., J.W. wrote the manuscript.

## Data and Code Availability

Raw and processed data are available at GEO under accession GSE313074. We have also uploaded all processed files to Zenodo (DOI: 10.5281/zenodo.17467446). All custom code used for genomic data analysis is available on our GitHub page: https://github.com/thewonlab/C9orf72-ALS.

