## Supplementary Figures for "Repeat expansions in *C9orf72* rewire the 3D chromatin landscape in ALS"

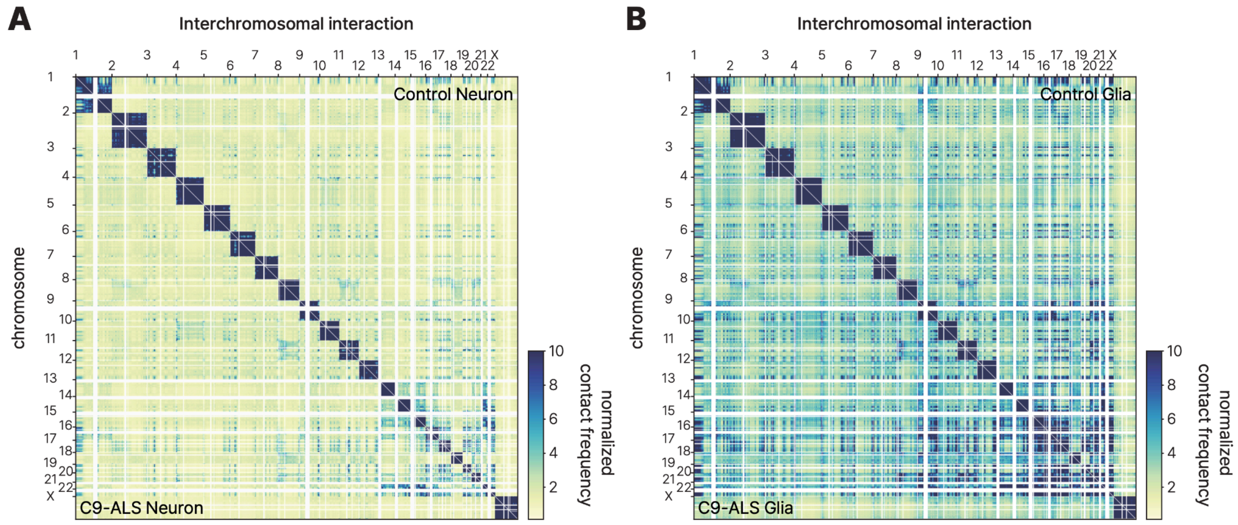


**Figure S1. Interchromosomal interactions in C9-ALS. A-B.** Interchromosomal interactions are largely preserved between C9-ALS and neurotypical controls in both neurons (**A**) and glia (**B**).

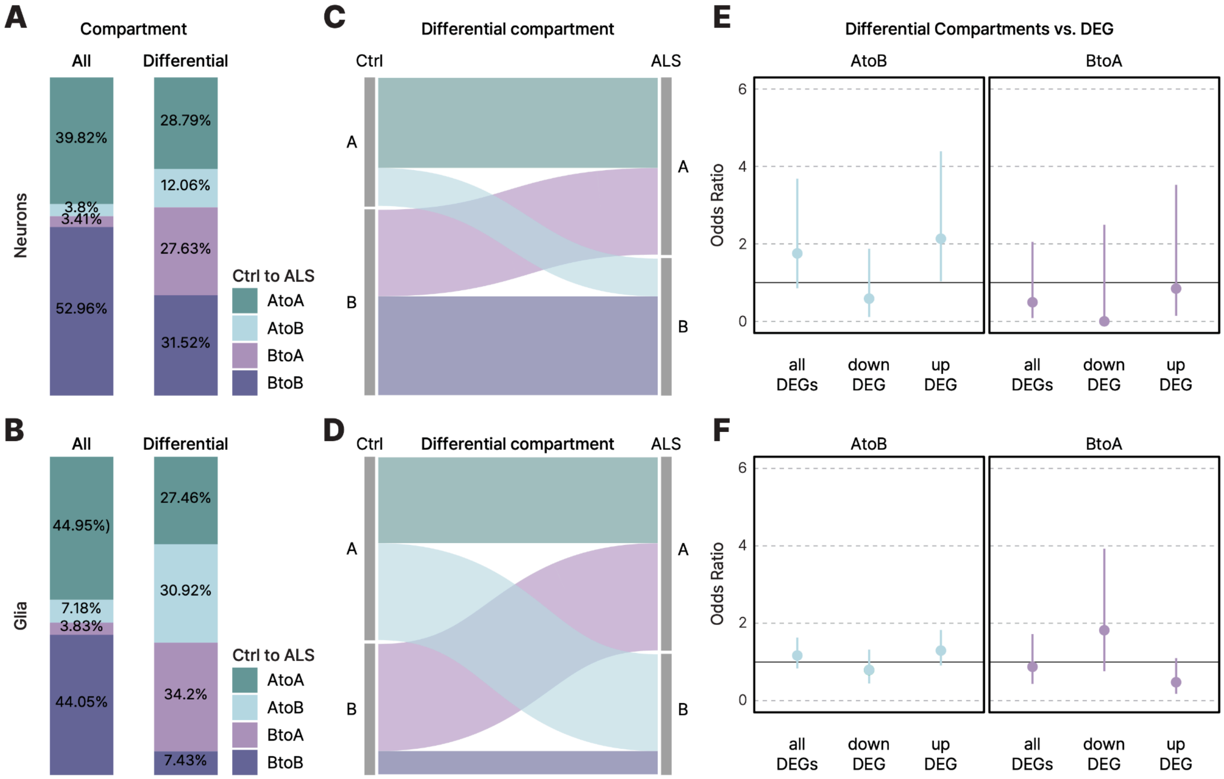


**Figure S2. A/B compartment organization in C9-ALS. A-B.** A/B compartment organization between C9-ALS and controls in neurons (**A**) and glia (**B**). **C-D.** Differential compartment profiles between C9-ALS and controls in neurons (**C**) and glia (**D**). **E-F.** Compartment switching shows no clear association with DEGs in C9-ALS neurons (**E**) and glia (**F**).

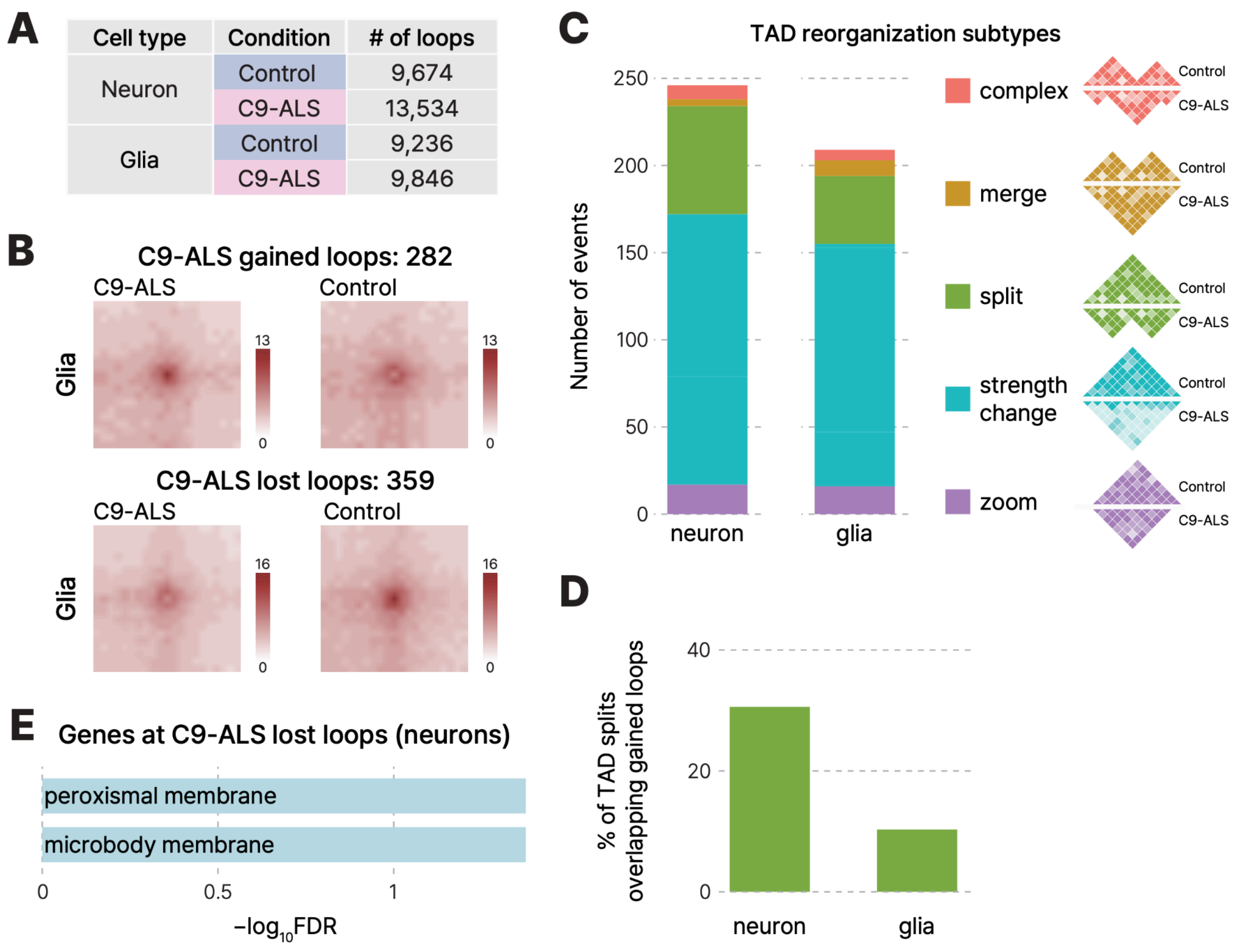


**Figure S3. Cell type-specific loops and TADs in C9-ALS. A.** Number of chromatin loops detected in each condition. **B.** Aggregated Peak Analysis (APA) results for C9-ALS gained loops and C9-ALS lost loops in glia. **C.** TAD reorganization in C9-ALS neurons and glia. **D.** C9-ALS gained loops frequently coincide with split TAD boundaries in neurons, but not in glia. **E.** Gene ontology analysis of genes anchored at C9-ALS lost loops.

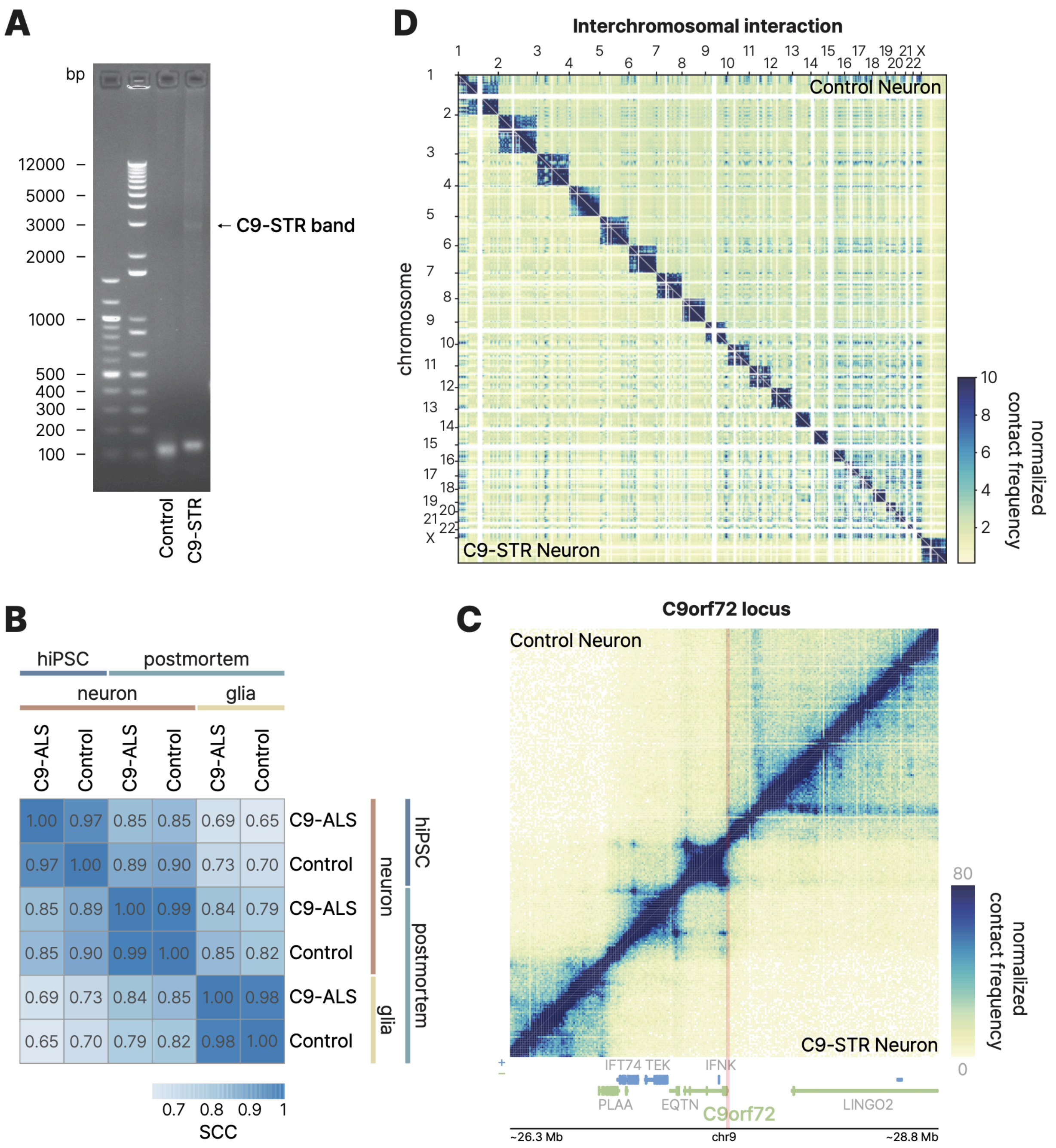


**Figure S4. Local and global chromosome conformation of C9-STR neurons. A.** Repeat PCR shows a ~3kb C9-STR expansion band in C9-STR hiPSCs, which is not present in isogenic control hiPSCs. **B.** Stratum-adjusted correlation coefficients (SCC) comparing postmortem neuronal and glial Hi-C libraries with hiPSC-derived neuronal Micro-C libraries. **C.** Local chromatin interactions at the *C9orf72* locus in hiPSC-derived neurons carrying the C9-STR expansion (bottom right) and isogenic controls (top left). **D.** Interchromosomal interactions in C9-STR neurons (bottom left) and isogenic controls (top right).

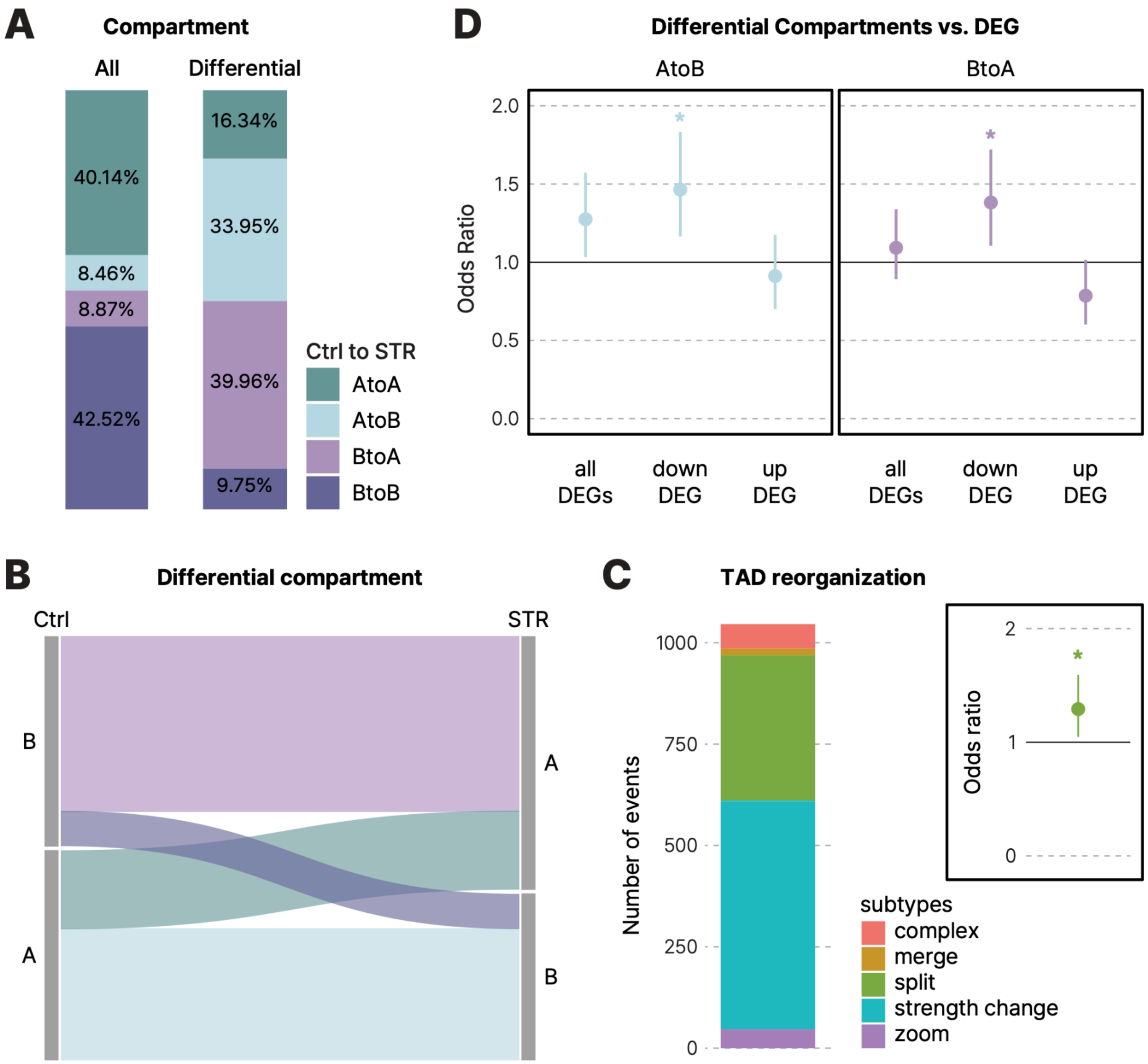


**Figure S5. A/B compartment organization associated with C9-STR expansion. A.** A/B compartment organization between C9-STR neurons and isogenic controls. **B.** Differential compartment profiles between C9-STR neurons and isogenic controls. **C.** TAD reorganization in C9-STR neurons. TAD splitting events overlapped more frequently with C9-STR gained loops than with stable loops. Two-sided Fisher's exact test, *P<0.05. **D.** Regions undergoing B-to-A compartment switching are enriched for downregulated genes in C9-STR neurons. Two-sided Fisher's exact test, *P<0.05.

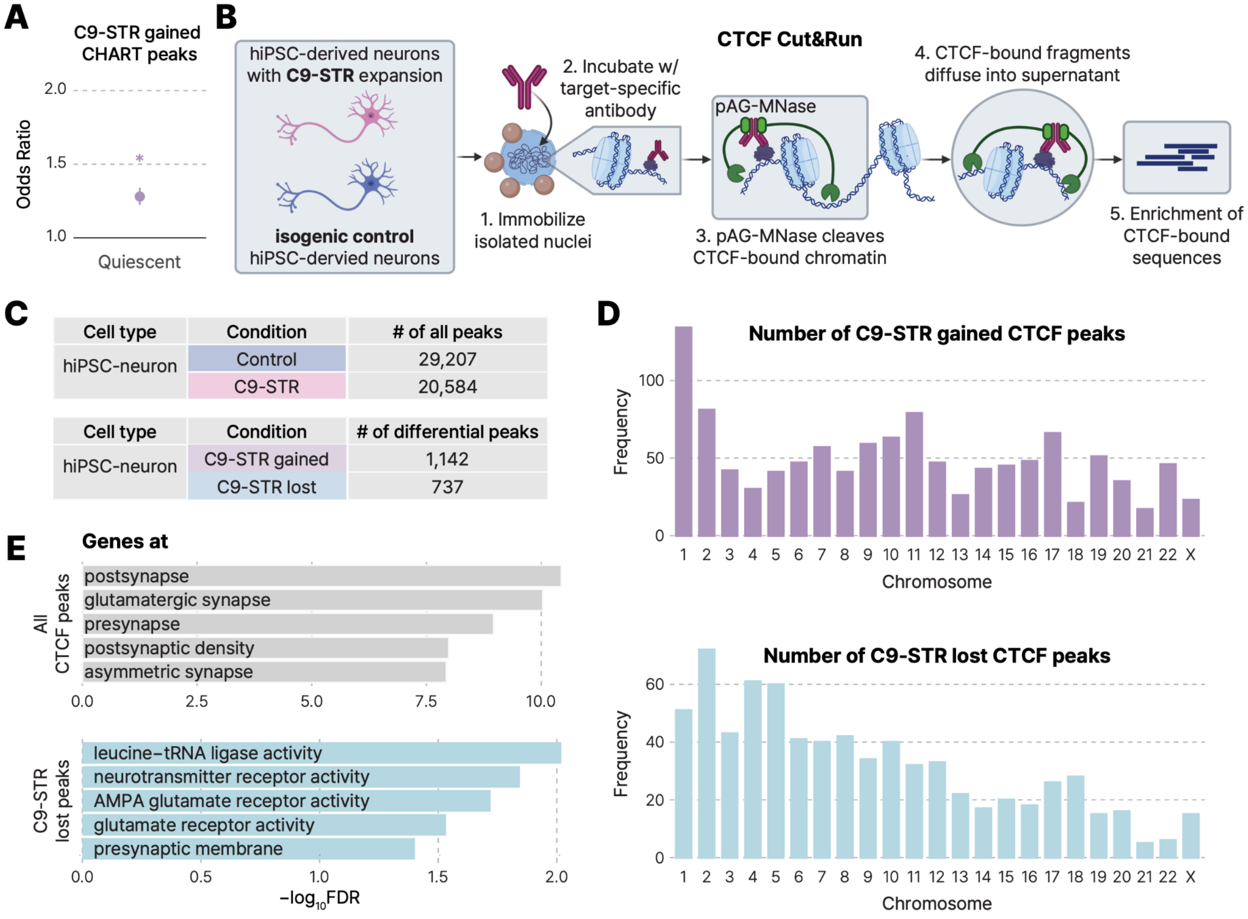


**Figure S6. Relationship between C9-STR gained loops and CTCF binding sites. A.** C9-ALS gained CHART peaks were enriched in quiescent chromatin states compared to the CHART input background. Two-sided Fisher exact test, *$p<2.2\times10^{-16}$. **B.** CTCF CUT&RUN was performed in hiPSC-derived neurons with C9-STR expansion and isogenic controls to profile the CTCF binding landscape. **C.** Total number of CTCF peaks in hiPSC-derived neurons with C9-STR expansion and isogenic controls (top), and the number of differential CTCF peaks gained or lost in C9-STR neurons (bottom). **D.** Genomic distribution of C9-STR gained (top) and lost (bottom) CTCF peaks. **E.** Gene ontology analysis of genes anchored at all CTCF peaks (top) and C9-STR lost CTCF peaks (bottom). No significant gene ontology enrichment was observed for genes anchored at C9-STR gained CTCF peaks.

### Supplementary Tables

**Table S1.** Compartment changes in C9-ALS glia, C9-ALS neuron, and C9-STR neuron

**Table S2.** Differential loops in C9-ALS glia, C9-ALS neuron, and C9-STR neuron

**Table S3.** Stable (non-differential) and differential TADs in C9-ALS glia, C9-ALS neuron, and C9-STR neuron

**Table S4.** Upregulated and downregulated DEGs between C9-STR and control neurons

**Table S5.** C9-STR gained CHART peaks

**Table S6.** De novo HOMER results for C9-STR gained CHART peaks

**Table S7.** C9-STR gained and lost CTCF peaks
